# Annexin A6 mediates calcium-dependent secretion of exosomes

**DOI:** 10.1101/2022.12.28.522125

**Authors:** Justin Krish Williams, Jordan Matthew Ngo, Randy Schekman

## Abstract

Exosomes are a subtype of extracellular vesicles (EV) that are secreted upon fusion of multivesicular bodies (MVBs) with the plasma membrane. In order to identify proteins responsible for this fusion event, we developed sensitive cellular and cell-free assays to measure exosome secretion. Our results suggest that exosome secretion is coupled to Ca^2+^-dependent plasma membrane repair. Using a targeted proteomics approach, we identify annexin A6 (ANXA6), a well-known plasma membrane repair protein, as a MVB binding protein and demonstrate that it is required for Ca^2+^-dependent exosome secretion both in intact and in permeabilized cells. Our results suggest that cells employ MVBs as a membrane source for plasma membrane lesion repair during physiological mechanical stress or treatment with a bacterial pore-forming toxin, and that this repair process results in the coincident secretion of exosomes. We propose that this unconventional secretion process may contribute significantly to the heterogeneity of EVs in biological fluids.

## Introduction

Protein secretion is an essential process that prokaryotes and eukaryotes utilize for intercellular communication. In eukaryotic cells, most secretory proteins are exported by the conventional secretory pathway which consists of co-translational signal peptide recognition by the signal recognition particle, polypeptide translocation through the Sec61 translocon, and anterograde transport from the endoplasmic reticulum to the Golgi apparatus via COPII vesicles (1–3). However, eukaryotic cells have also developed alternative secretory processes that bypass the conventional pathway (4). Some well-documented examples include the secretion of leaderless cargoes (proteins that lack a signal peptide) such as interleukin-1β, and the egress of cytoplasmic proteins and RNAs within extracellular vesicles (EVs) (5–7).

EVs are membrane-enclosed compartments that are exported to the extracellular milieu of cells in culture and in vivo. Eukaryotic cells secrete EV subpopulations that can be classified broadly into two major categories with distinct biogenesis pathways: microvesicles and exosomes (6, 8). Microvesicles consist of a heterogeneous pool of EVs that form by outward budding from the plasma membrane (9, 10). Exosomes are 30-150 nm vesicles formed by inward budding of the limiting membrane of late endosomes to produce multivesicular bodies (MVBs) that contain intraluminal vesicles (ILVs). Upon fusion of MVBs with the plasma membrane, ILVs are released to the extracellular space as exosomes (11). Exosomes have potential utility as biomarkers for disease because they are enriched with small non-coding RNA species such as miRNAs, tRNAs, and yRNAs (12–14). Additionally, elevation of cytosolic Ca^2+^ with ionophores induces rapid EV secretion in a Rab27a, Munc13-4-dependent manner (15, 16). Despite these reports, how this elevation might occur physiologically, and which other genes are involved, remain uncertain.

Like MVBs, lysosomes can fuse at the plasma membrane to release their luminal content. Lysosome exocytosis is a response to plasma membrane rupture (17). Disruption of the plasma membrane results in the influx of extracellular Ca^2+^ into the cytoplasm, triggering a repair cascade that includes lysosome exocytosis. Some reports have suggested that, unlike lysosomes, MVBs do not undergo Ca^2+^-dependent exocytosis (18, 19). Despite the similarities between the two organelles, it is unclear if MVBs, like lysosomes, fuse with the plasma membrane upon Ca^2+^ influx and participate in plasma membrane repair.

To determine if MVBs and lysosomes utilize a common exocytic pathway and if both participate in plasma membrane repair, we investigated the release of exosomes upon elevation of cytosolic Ca^2+^. First, we developed an improved nanoluciferase (Nluc) reporter system to quantify exosome secretion that is sensitive, linear, and amenable to high throughput. Using this assay, we establish that a Ca^2+^ ionophore, a pore-forming cytolysin, and physiological mechanical stress all stimulate exosome secretion in an extracellular Ca^2+^-dependent manner. We then identified annexin A6 as a Ca^2+^-dependent MVB binding protein that is required for Ca^2+^-dependent exosome secretion. Next, we developed a streptolysin O (SLO)-permeabilized cell reaction that recapitulates Ca^2+^- and ANAX6-dependent exosome secretion. Finally, we demonstrate that the two annexin domains of ANXA6 become enriched at different membranes upon elevation of cytosolic Ca^2+^. Our results demonstrate that MVBs undergo ANXA6-dependent exocytosis upon plasma membrane damage, and that this process is coupled to exosome secretion.

## Results

### Design and validation of an endogenous CD63-Nluc exosome secretion assay

We sought to develop a cell-based exosome secretion assay that is quantitative, sensitive, amenable to high-throughput, and able to distinguish cell debris and *bona fide* exosomes. A previous study developed a luminescence-based assay to quantify exosome secretion by inserting Nluc into the endogenous locus of the tetraspanin protein CD63 (20). We obtained the HCT116 CD63Nluc-KI #17 cell line (referred to herein as CD63-Nluc cells) from this group and built upon their assay.

Internalization of CD63 during ILV biogenesis results in the amino- and carboxyl-termini oriented to the exosome lumen. We leveraged this topology along with the relative membrane permeabilities of the substrate and an inhibitor of Nluc to develop a quantitative cellular assay to monitor the secretion of exosomes (Fig. 1*A*). Furimazine, the substrate of Nluc, permeates through cellular membranes and enables luminescence from both cellular debris and intact CD63-Nluc exosomes present within the conditioned medium of cultured CD63-Nluc cells. The addition of a membrane-impermeable Nluc inhibitor eliminates luminescence derived from cellular debris where Nluc is readily accessible, while not affecting the luminescence derived from intact CD63-Nluc exosomes incubated with furimazine (21). Finally, solubilization of cellular membranes by addition of the non-ionic detergent Triton X-100 (TX-100) exposes all CD63-Nluc to the inhibitor. Addition of the membrane-impermeable Nluc inhibitor depleted a significant portion of luminescence derived from the conditioned medium of CD63-Nluc cells (∼30%) and membrane solubilization by TX-100 reduced luminescence to background levels (Fig. 1*B*).

**Fig. 1.**
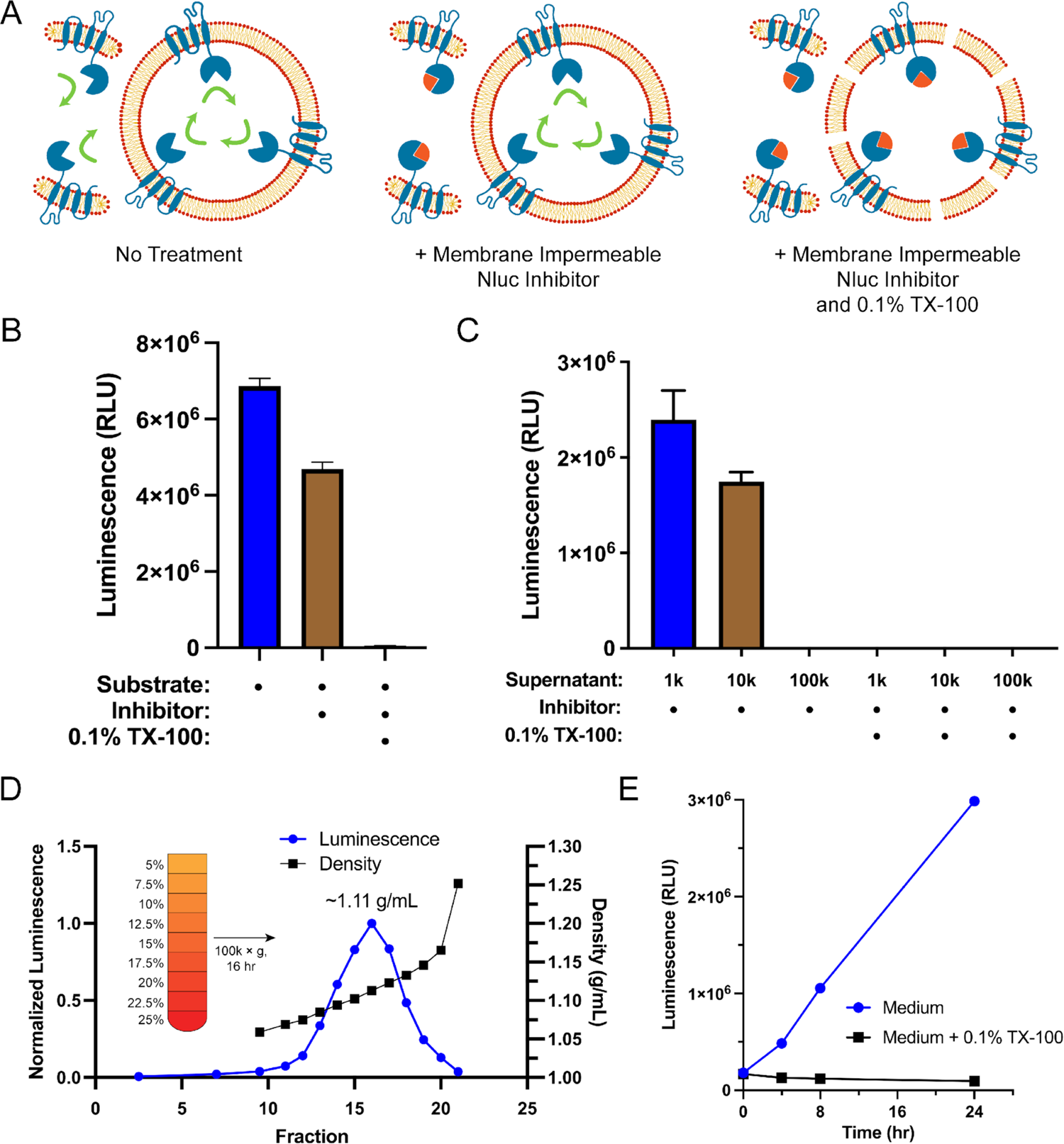
Endogenous CD63-Nluc is a faithful reporter of exosome secretion. (*A*) Schematic illustrating the cellular CD63-Nluc secretion assay. (*B*) Luminescence from the conditioned medium of CD63-Nluc cells with or without furimazine, the membrane-impermeable Nluc inhibitor, and 0.1% TX-100 are shown. (*C*) Luminescence derived from the supernatant fraction of CD63-Nluc conditioned medium subjected to differential centrifugation (1k, 10k, and 100k) with or without 0.1% TX-100 are shown. (*D*) Membrane-protected luminescence (in blue circles) and buoyant density (in black squares) of CD63-Nluc conditioned medium subjected to a high-resolution linear density gradient are shown. (*E*) Membrane-protected luminescence from CD63-Nluc conditioned medium collected over 24 h with (blue circles) or without 0.1% TX-100 (black squares). Data plotted represent the means of 3 independent experiments and error bars represent standard deviations. Note, for Fig. 1*D* and 1*E*, the error bars are smaller than the dots in the image.

To further validate the localization of the CD63-Nluc luminescence signal to sedimentable vesicles, we subjected the conditioned medium fraction to differential centrifugation and found that CD63-Nluc luminescence was removed after high-speed sedimentation or upon the addition of TX-100 (Fig. 1*C*). Next, we obtained a partially-clarified conditioned medium fraction by differential centrifugation and fractionated the material on a linear iodixanol gradient (Fig. 1*D*). CD63-Nluc vesicles equilibrated at a density of 1.11 g/ml, a buoyancy similar to published reports (22). In a time course over 24 h, we measured secreted CD63-Nluc luminescence in the presence of Nluc inhibitor alone or with TX-100 and found the signal increased linearly, consistent with exosome secretion over time (Fig. 1*E*). The luminescence signal in the presence of the Nluc inhibitor and TX-100 remained constant, consistent with cell rupture/detachment during medium change. Thus, our modified CD63-Nluc assay faithfully reported the secretion of *bona fide* CD63-positive exosomes and we have used this method to assess the molecular requirements for exosome secretion.

### Elevation of cytosolic Ca^2+^ stimulates exosome secretion

We investigated the relationship between elevation of cytosolic Ca^2+^ levels and exosome secretion. In agreement with previous reports (15), influx of Ca^2+^ into the cytoplasm using the Ca^2+^ ionophore, ionomycin (5 µM, 30 min), induced rapid, robust exosome secretion compared to the level of secretion of the vehicle control (Fig. 2*A*).

**Fig. 2.**
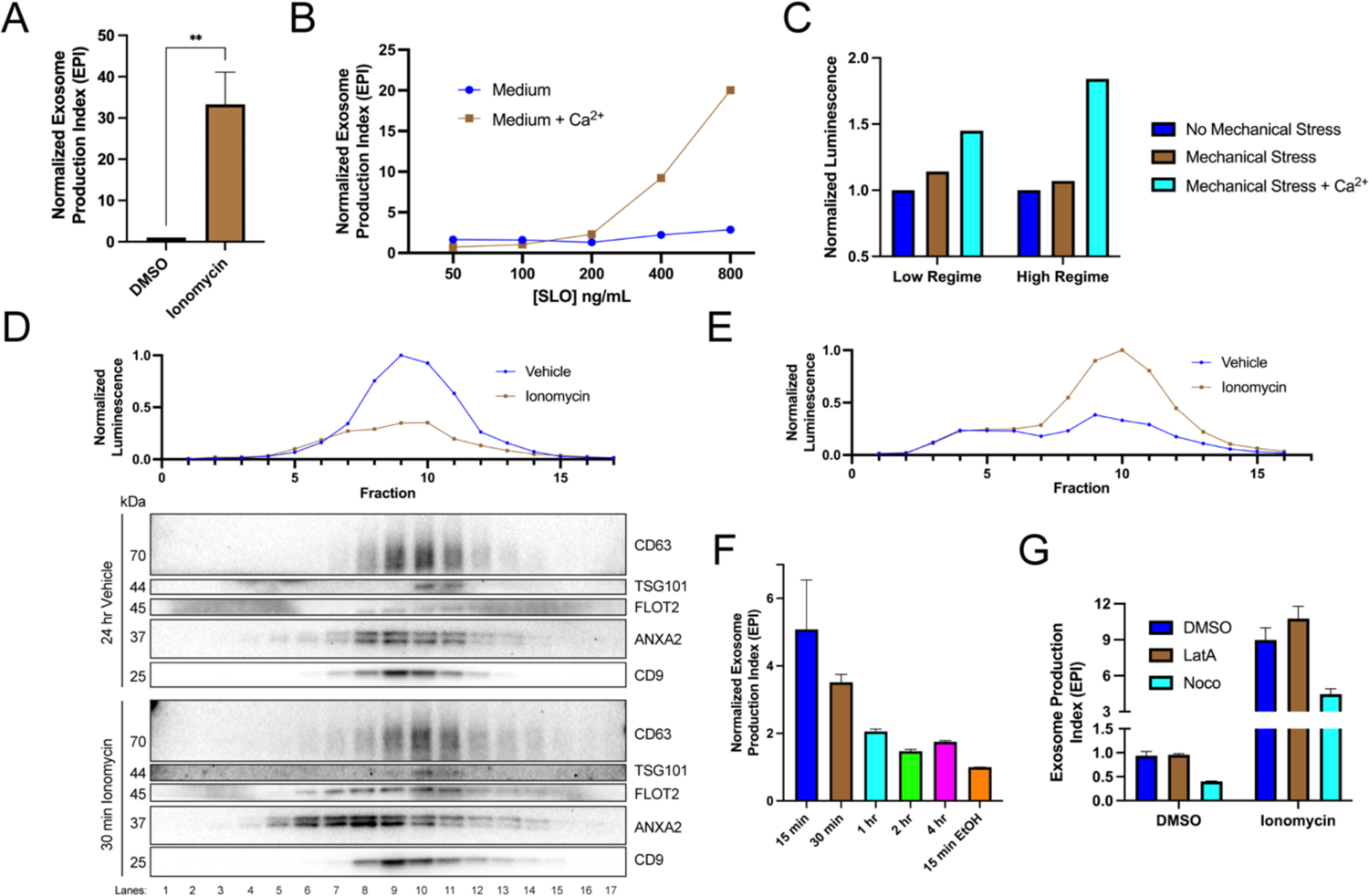
Elevation of cytosolic Ca^2+^ levels promotes exosome secretion. (*A*) Normalized exosome production from CD63-Nluc cells treated with 5 µM ionomycin or DMSO (ionomycin vehicle) are shown. (*B*) Normalized exosome production from CD63-Nluc cells treated for 30 min with increasing concentrations of SLO, with or without 1.8 mM extracellular Ca^2+^. Note, error bars are smaller than the dots in the image. (*C*) Normalized luminescence derived from CD63-Nluc cells treated with a high or low dose of mechanical stress, with or without 1.8 mM extracellular Ca^2+^. (*D*) Iodixanol gradient fractionation of conditioned medium 100k x g pellet fraction is shown. Conditioned medium from cells treated for 30 min with 5 µM ionomycin or 24 h vehicle are compared. Line graphs show distribution of CD63-Nluc luminescence (with membrane-impermeable inhibitor added) across the linear gradient. Immunoblots show distribution of several EV markers across the linear gradient. (*E*) Iodixanol gradient fractionation of conditioned medium 10k x g supernatant fractions are shown. Conditioned medium from cells treated for 4 h with 5 µM ionomycin or 4 h vehicle are compared. Line graphs show distribution of CD63-Nluc luminescence (with membrane-impermeable inhibitor added) across the linear gradient. (*F*) Relative rate of exosome production over time is shown. CD63-Nluc cells were treated with ionomycin or EtOH vehicle where indicated and the medium was replaced for the last 15 min of each timepoint. The replaced medium was used to calculate the exosome production index. (*G*) Normalized exosome production from 30 min of 5 µM ionomycin or DMSO vehicle, co-treated with DMSO vehicle, 1 µM latrunculin A (LatA), or 10 µM nocodazole (Noco) is shown. Data plotted represent the means from 3 independent experiments and error bars represent standard deviations. Statistical significance was performed using a Student’s T-test (**p<0.01).

During plasma membrane repair, caused by plasma membrane disruption (17, 23–25) or simulated with Ca^2+^ ionophore treatment (18), an influx of extracellular Ca^2+^ triggers lysosome exocytosis. We reasoned that MVBs, like lysosomes, might fuse with the plasma membrane to facilitate membrane repair. Upon treatment with the pore-forming toxin, SLO, or application of a physiological mechanical force, exosomes were secreted in a Ca^2+^-dependent manner (Fig. 2*B* and *C*). Treatment with 400 ng/ml or 800 ng/ml of SLO significantly increased exosome production, dependent on the presence of extracellular CaCl_2_ (1.8 mM) (Fig. 2*B*). We noted that SLO permeabilized exosomes as well as cells, as SLO treatment of conditioned medium decreased the CD63-Nluc signal when the membrane-impermeable Nluc inhibitor was present (Fig. S1*A*). Thus, our assay likely underestimates SLO-induced exosome secretion. For mechanical stress experiments, cells were pumped slowly through a narrow-gauge needle to simulate a capillary (Fig. 2*C*). In the low mechanical stress regime, cells transiently experienced ∼89 dyn/cm^2^ maximum fluid shear stress, while in the high mechanical stress regime, cells transiently experienced ∼178 dyn/cm^2^ maximum fluid shear stress (26). This form of mechanical stress increased exosome production, but only when CaCl_2_ was present in the conditioned medium. Endothelium and circulating lymphocytes routinely experience mechanical stress of ∼95 dyn/cm^2^ and transiently experience up to ∼3000 dyn/cm^2^ (26).

Next, we used linear iodixanol density gradient fractionation to assess if the Ca^2+^-dependent increase in extracellular CD63 was attributable to exosomes as opposed to the secretion of intact endosomes contained within plasma membrane-derived vesicles (22). EVs from cells treated with ionomycin for 30 min or vehicle for 24 h were separated on a high-resolution linear iodixanol gradient. A 30 min ionomycin treatment induced the secretion of low buoyant density, ANXA2- and FLOT2-positive vesicles (Fig. 2*D*, Fraction #8) compared to the 24 h vehicle control. ANXA2 is a marker for a subpopulation of plasma membrane-derived EVs (22). Alternatively, CD63 and Nluc signals equilibrated to higher buoyant density fractions corresponding to the position expected for exosomes (Fig. 2*D*, Fraction #10). A comparison of ionomycin- and vehicle-treated samples showed drug treatment increased the Nluc luminescence in putative exosome fractions (Fig. 2*E*, Fraction #10). Treatment of cells with ionomycin for 4 h did not induce apoptotic cell death (Fig. S1*B*) (27).

MVBs traffic toward the cell periphery on microtubules or actin filaments (28). Ionomycin treatment elevated the rate of exosome secretion over vehicle control even 4 h after initiation of the treatment (Fig. 2*F*). This result suggested that MVBs may continuously traffic toward and fuse with the plasma membrane. We tested this model using cytoskeleton depolymerizing drugs. Treatment with nocodazole, a microtubule polymerization inhibitor, but not latrunculin A, an actin polymerization inhibitor, reduced vehicle- and ionomycin-induced exosome secretion (Fig. 2*G*, Fig. S1*C* and *D*). Ionomycin also partially redistributed Lamp1-positive compartments toward the cell periphery (Fig. S1*E*).

We conclude that the influx of extracellular Ca^2+^ caused by several inducers of plasma membrane lesions triggers exosome secretion, and that sustained exosome secretion is dependent on the anterograde trafficking of MVBs on microtubules.

### Identification of candidate proteins involved in Ca^2+^-dependent exosome secretion

Next, we sought to identify cytosolic proteins that are recruited in a Ca^2+^-dependent manner to MVBs. Such proteins may be involved in Ca^2+^-dependent exosome secretion and plasma membrane repair. CD63-Nluc cells were ruptured by homogenization and a post nuclear supernatant fraction was supplemented with 1mM CaCl_2_ or 1 mM EGTA and mixed with immobilized Nluc antibody in order to immunoprecipitate MVBs and associated peripheral membrane proteins. After washing the immunoprecipitated MVBs to remove other organelles, Ca^2+^-dependent MVB binding proteins were eluted from the IP pellet fraction using 2 mM EGTA. MVB proteins retained after the EGTA elution were solubilized using 0.2% TX-100 (Fig. 3*A*). The 0.2% TX-100 elution fractions were significantly enriched in LAMP1 and diminished in GAPDH compared to the input, regardless of the presence of CaCl_2_ or EGTA in the elution (Fig. 3*B*). A gel stained for total protein showed four intense protein bands unique to the EGTA-eluted sample (Fig. 3*C*). These proteins were absent when the Nluc antibody was omitted or when the post-nuclear supernatant was treated with 1 mM EGTA instead of 1 mM CaCl_2_. Three sections of the gel were excised and analyzed by mass spectrometry (Fig. 3*D*). We identified annexin A6 (ANXA6) and copine 3 (CPNE3) in the upper gel slice, annexin A2 (ANXA2) in the middle gel slice, and S100 calcium-binding protein A10 (S100A10) in the lower gel slice.

**Fig. 3.**
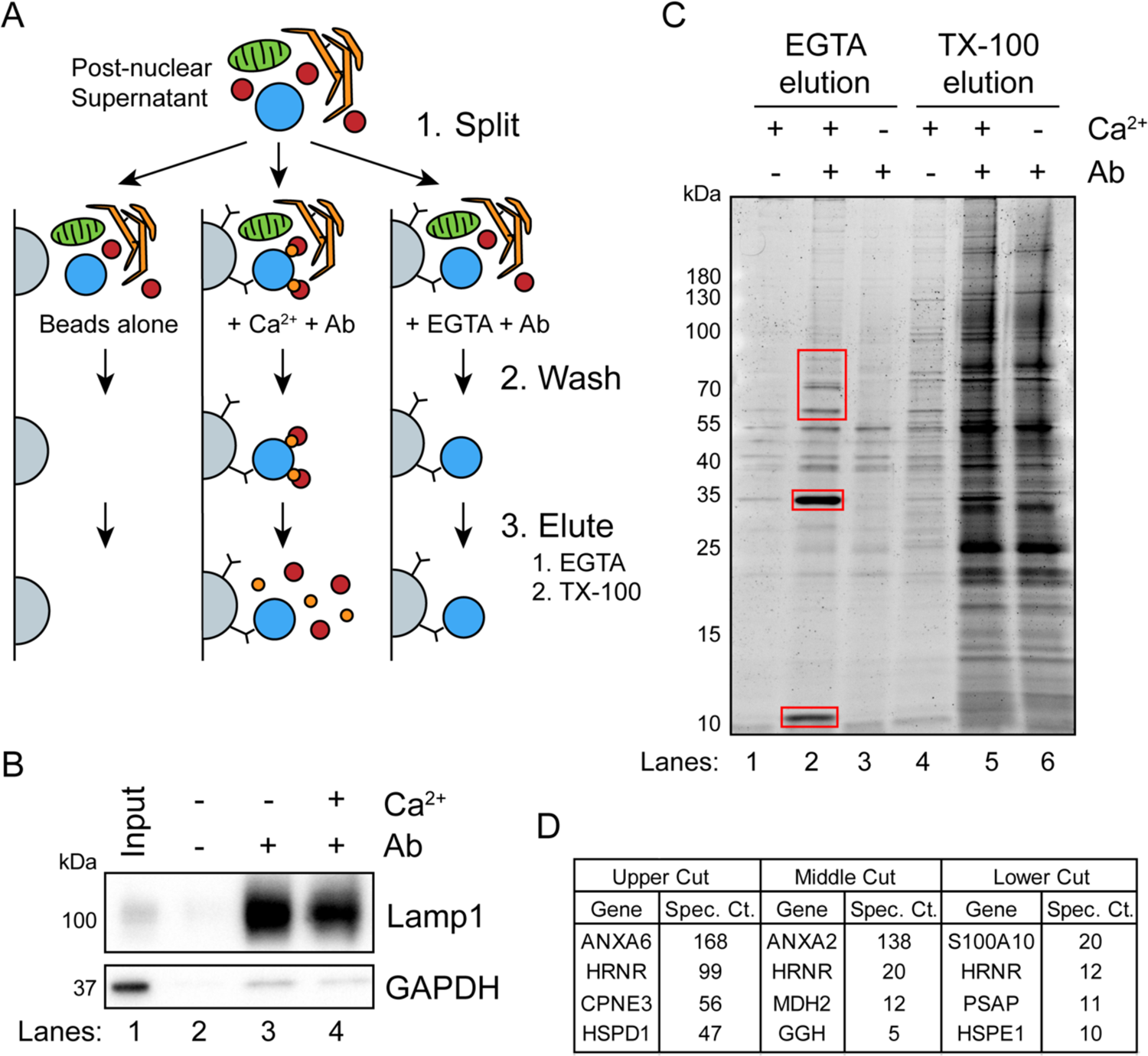
A targeted proteomics approach identifies genes important for Ca^2+^-dependent exosome secretion. (*A*) Schematic illustrating the pulldown of Ca^2+^-dependent MVB/lysosome binding proteins (Grey-Beads, Blue-MVB, Green-Mitochondria, Orange-ER, Red-proteins, Gold-calcium). (*B*) Immunoblot analysis of LAMP1 and GAPDH from the TX-100 elutions, relative to the input, are shown. (*C*) Total protein gel (Sypro Ruby stained) of eluted fractions is shown. Red boxes indicate gel cuts sent for proteomic analysis. (*D*) Table depicting the top 4 proteomic hits from each gel cut are shown, excluding keratin family proteins.

### Annexin A6 is required for Ca^2+^-dependent exosome secretion

We tested the effect of knocking-down genes encoding the Ca-dependent MVB-binding proteins identified in our proteomic analysis. ANXA6 and CPNE3, but not ANXA2, were efficiently knocked-down with two different shRNAs (Fig. 4*A* and *B*), and in each case, Ca^2+^-dependent exosome secretion decreased relative to a GFP knockdown control (Fig. 4*C* and *D*). One of the ANXA6 shRNAs also slightly decreased constitutive exosome secretion relative to the GFP control, although the effect was relatively small (Fig. 4*C*). A polyclonal knockout of ANXA6 also decreased exosome secretion relative to a non-targeting control (Fig. S2*A* and *B*).

**Fig. 4.**
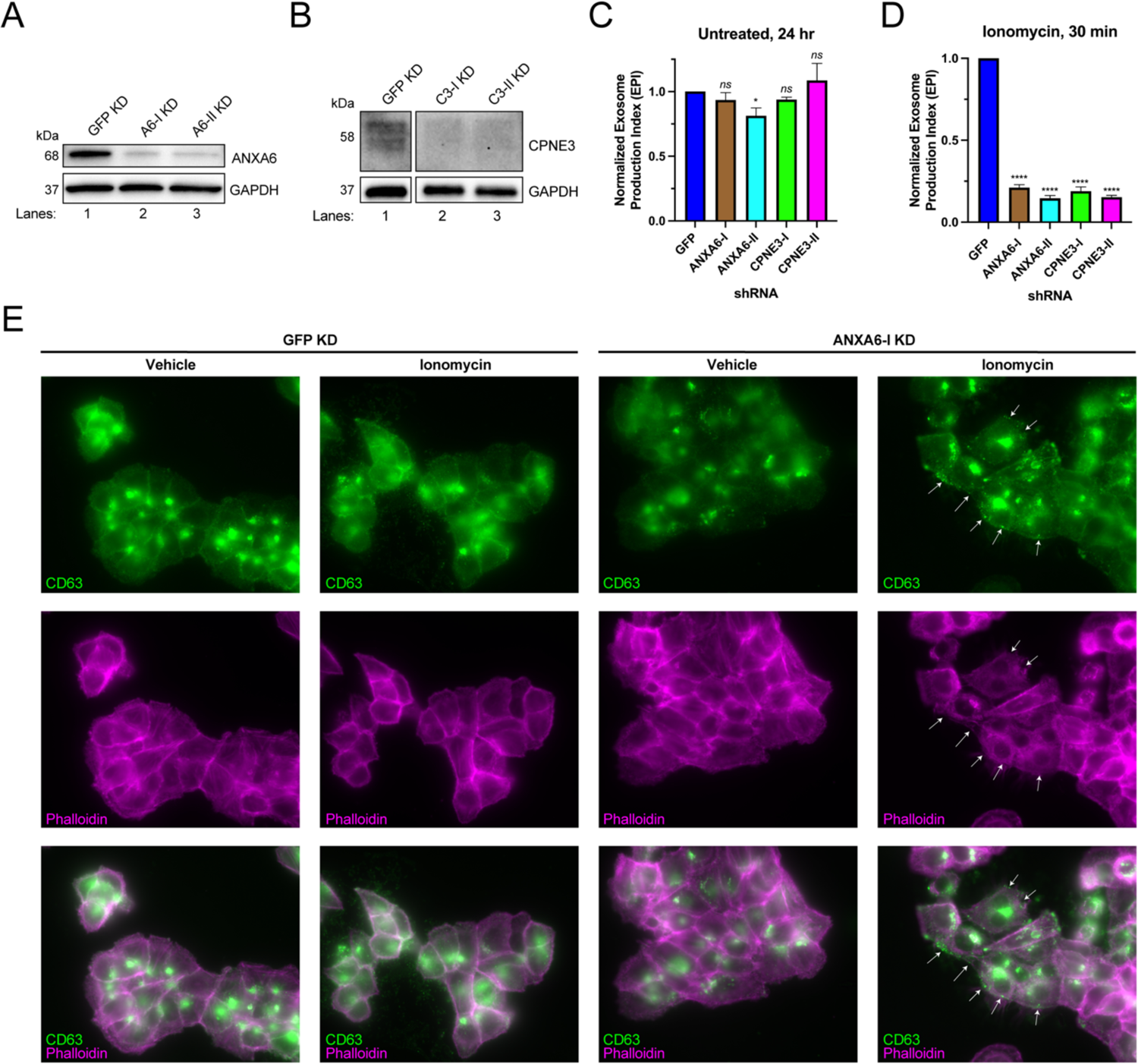
ANXA6 depletion blocks ionomycin-mediated exosome secretion. (*A*) Immunoblot analysis of ANXA6 and GAPDH expression from GFP, ANXA6-I, and ANXA6-II shRNA CD63-Nluc cells are shown. (*B*) Immunoblot analysis of CPNE3 and GAPDH expression from GFP, CPNE3-I, and CPNE3-II shRNA CD63-Nluc cells are shown. (*C*) Normalized exosome production derived from GFP, ANXA6-I, ANXA6-II, CPNE3-I, CPNE3-II shRNA CD63-Nluc cells grown in conditioned medium for 24 h are shown. (*D*) Normalized exosome production derived from GFP, ANXA6-I, ANXA6-II, CPNE3-I, CPNE3-II shRNA CD63-Nluc cells treated with ionomycin for 30 min are shown. (*E*) CD63 immunofluorescence and phalloidin staining of GFP or ANXA6-I shRNA CD63-Nluc cells after 30 min of DMSO or ionomycin treatment is shown. White arrows indicate peripheral CD63 puncta. Data plotted represent the means from 3 independent experiments and error bars represent standard deviations. Statistical significance was performed using an ANOVA (*p<0.05, ****p<0.0001, and ns = not significant).

In order to assess how the knock-down of ANXA6 affected exosome secretion, we visualized CD63-positive compartments after Ca^2+^ influx. CD63-Nluc cells were treated with ionomycin or vehicle control for 30 min, fixed, and analyzed using immunofluorescence microscopy. We observed an accumulation of CD63-positive vesicles at the cell periphery, particularly in ionomycin-treated ANXA6 knockdown cells (Fig. 4*E*). We suggest that ANXA6 may be necessary for Ca^2+^-dependent exosome secretion, possibly at the point of fusion between the MVB and the plasma membrane.

### Cell-free reconstitution of Ca^2+^- and ANAX6-dependent exosome secretion

Many studies probing genes that may contribute to exosome secretion have relied on depletion or overexpression experiments in live cells. We sought to develop a cell-free assay to allow a direct assessment of the roles of different gene products in exosome secretion.

SLO is a potent bacterial toxin secreted by group A streptococci that forms stable pores within cholesterol-containing biological membranes in a temperature-dependent manner (29). SLO has been used to permeabilize cells to reconstitute various intracellular membrane trafficking and organelle exocytosis reactions because it forms stable pores that are large enough to allow the diffusion of large proteins, but not intact organelles, across the plasma membrane (30–32). We leveraged the cholesterol- and temperature-dependent properties of SLO to establish a cell-free reaction that would allow us to study MVB exocytosis and coincident exosome secretion.

Reaction components including an ATP regeneration system (ATP^r^), GTP, cytosol, and Ca^2+^ were mixed with SLO-permeabilized cells and incubated at 30°C and evaluated for exosome secretion just as in the cell-based exosome secretion assay (Fig. 5*A*). We observed that addition of either rat liver cytosol or Ca^2+^ promoted exosome secretion slightly, and that the addition of both rat liver cytosol and Ca^2+^ promoted exosome secretion ∼12-fold over the control condition (Fig. 5*B*). The addition of HCT116 WT cytosol and Ca^2+^ promoted exosome secretion ∼15-fold over the control condition (Fig. 5*C*). To assess the energy dependence of this reaction, we repeated the experiment using nucleotide-depleted cytosol in either the absence or presence of the ATP^r^ and GTP (Fig. 5*D*). We observed that the Ca^2+^-only reaction was stimulated by the addition of the ATP^r^ and GTP (columns 3 and 7), whereas the stimulatory effect obtained with cytosol alone appeared to be nucleotide-independent (columns 2 and 6). The Ca^2+^ conditional and partial ATP dependence of the reaction may reflect distinct pools of MVBs, some perhaps already bound to the plasma membrane.

**Fig. 5.**
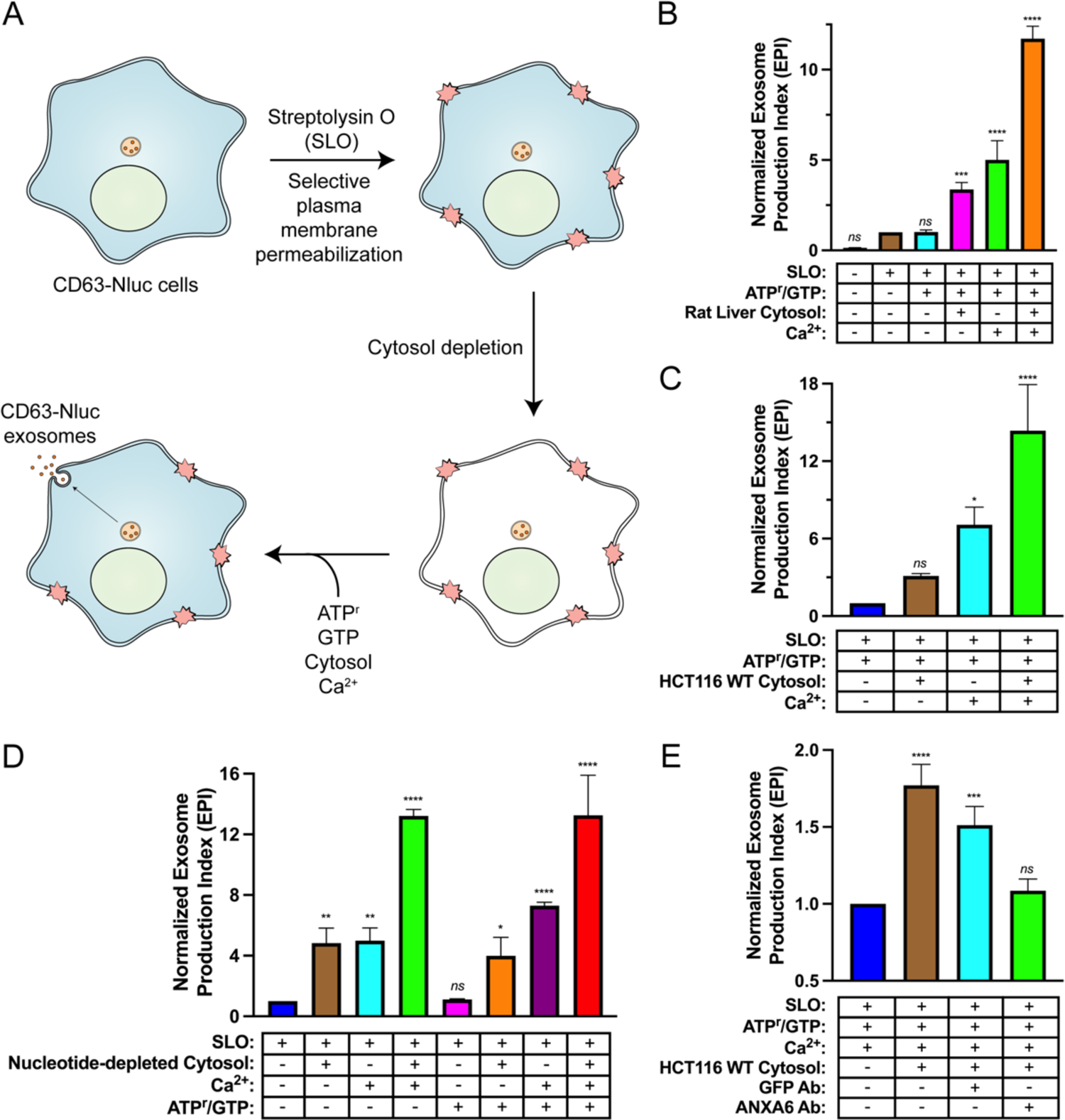
Cell-free reconstitution of Ca^2+^- and ANXA6-dependent exosome secretion. (*A*) Schematic illustrating the cell-free exosome secretion assay. (*B*) Cell-free exosome secretion reactions with or without SLO, ATP^r^/GTP, rat liver cytosol, and Ca^2+^ are indicated. (*C*) Cell-free exosome secretion assays with or without HCT116 WT cytosol and Ca^2+^ are shown. (*D*) ATP requirements for the cell-free exosome secretion assay. Reactions with or without nucleotide-depleted rat liver cytosol, Ca^2+^, and ATP^r^/GTP are indicated. (*E*) Requirement of ANXA6 in the cell-free exosome secretion assay. Reactions with or without HCT116 WT cytosol, an anti-GFP rabbit IgG antibody, and an anti-ANXA6 rabbit IgG antibody are depicted. Data plotted represent the means from 3 independent experiments and error bars represent standard deviations. Statistical significance was performed using an ANOVA (*p<0.05, **p<0.01, ***p<0.001, ****p<0.0001, and ns = not significant).

The participation of ANAX6 in the reconstitution was assessed with blocking IgG antibodies targeting epitopes on either GFP or ANAX6. Addition of an anti-GFP antibody slightly decreased exosome secretion whereas the equivalent addition of a knockout-validated anti-ANAX6 antibody decreased exosome secretion to a background level seen in a reaction without added cytosol (Fig. 5*E*). These results suggest a direct role of ANAX6 in the docking or fusion of MVBs at the cell surface.

### Annexin A6 truncations localize to different membranes upon ionomycin treatment

After demonstrating that ANXA6 is required for Ca^2+^-dependent exosome secretion in cells and in a cell-free reaction, we sought to probe the membrane recruitment of distinct domains of ANXA6 upon Ca^2+^ influx. Previous studies have demonstrated that ANXA6 is recruited to a peripheral “repair cap” that is formed at the site of plasma membrane lesions (33). Unlike other members of the annexin protein family, ANXA6 contains two complete annexin domains, ANXA6(N) and ANXA6(C). To probe the roles of these domains, we generated fluorescent ANXA6 full length (FL), ANXA6(N), and ANXA6(C) fusion constructs and probed their localization in U-2 OS cells by super-resolution microscopy.

We observed that ANXA6(FL), ANXA6(N) and ANXA6(C) all localized to the cytoplasm in unperturbed cells (Fig. 6*A*). However, upon addition of ionomycin, ANXA6(FL) was localized to the plasma membrane and to intracellular vesicles (Fig. 6*B*). We also observed the budding of plasma membrane-derived vesicles in the presence of ionomycin. These may correlate to the low buoyant density vesicles marked by ANXA2 and FLOT2 that were released from ionomycin-treated cells (Fig. 2*D*, Fraction #8). Finally, fluorescently-tagged ANXA6(N) and ANXA6(C) localized to distinct subcellular membranes after ionomycin treatment (Fig. 6*C*). Ca^2+^ stimulation relocalized ANXA6(N) to the cell periphery, whereas ANXA6(C) was observed on intracellular vesicles. The two domains may serve to dock MVBs to the plasma membrane.

**Fig. 6.**
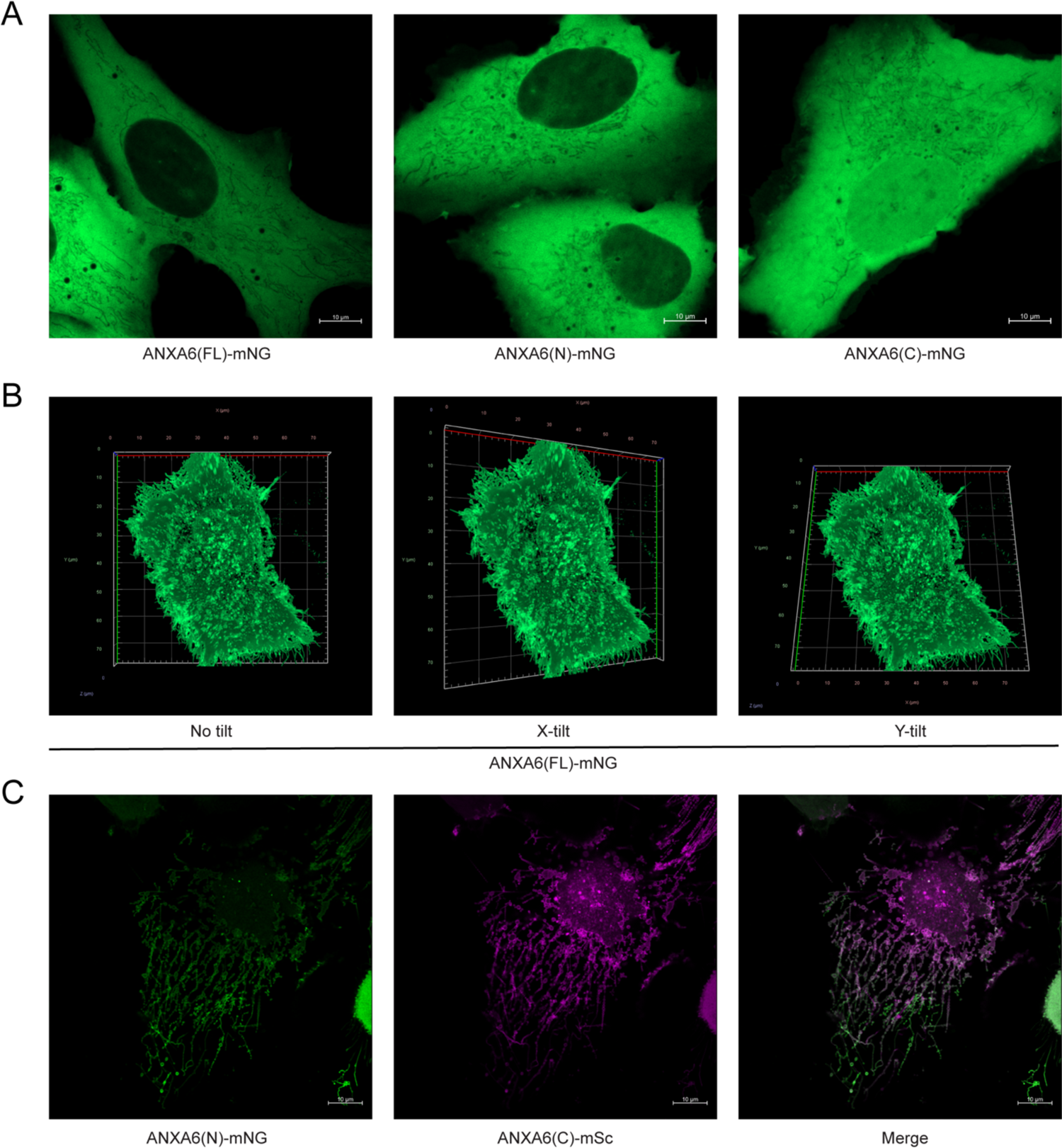
Localization of full-length and truncated ANXA6 constructs with or without ionomycin treatment. (*A*) Airyscan microscopy of U-2 OS cells expressing ANXA6(FL)-mNG (left), ANXA6(N)-mNG (middle), and ANXA6(C)-mNG (right). Scale bars: 10 µm. (*B*) Airyscan three-dimensional reconstruction tilt series of an ionomycin-treated U-2 OS cell expressing ANXA6(FL)-mNG. (*C*) Airyscan microscopy of an ionomycin-treated U-2 OS cell co-expressing ANXA6(N)-mNG (green) and ANXA6(C)-mSc (magenta). Scale bars: 10 µm.

## Discussion

Our results suggest that MVBs participate in Ca^2+^-dependent plasma membrane repair in HCT116 cells (Fig. 7). We established cellular and biochemical exosome secretion assays to recapitulate and interrogate this process. Targeted proteomics was used to identify ANXA6 as a Ca^2+^-dependent MVB-binding protein that stimulates Ca^2+^-dependent exosome secretion, in cells and a cell-free reaction.

**Fig. 7.**
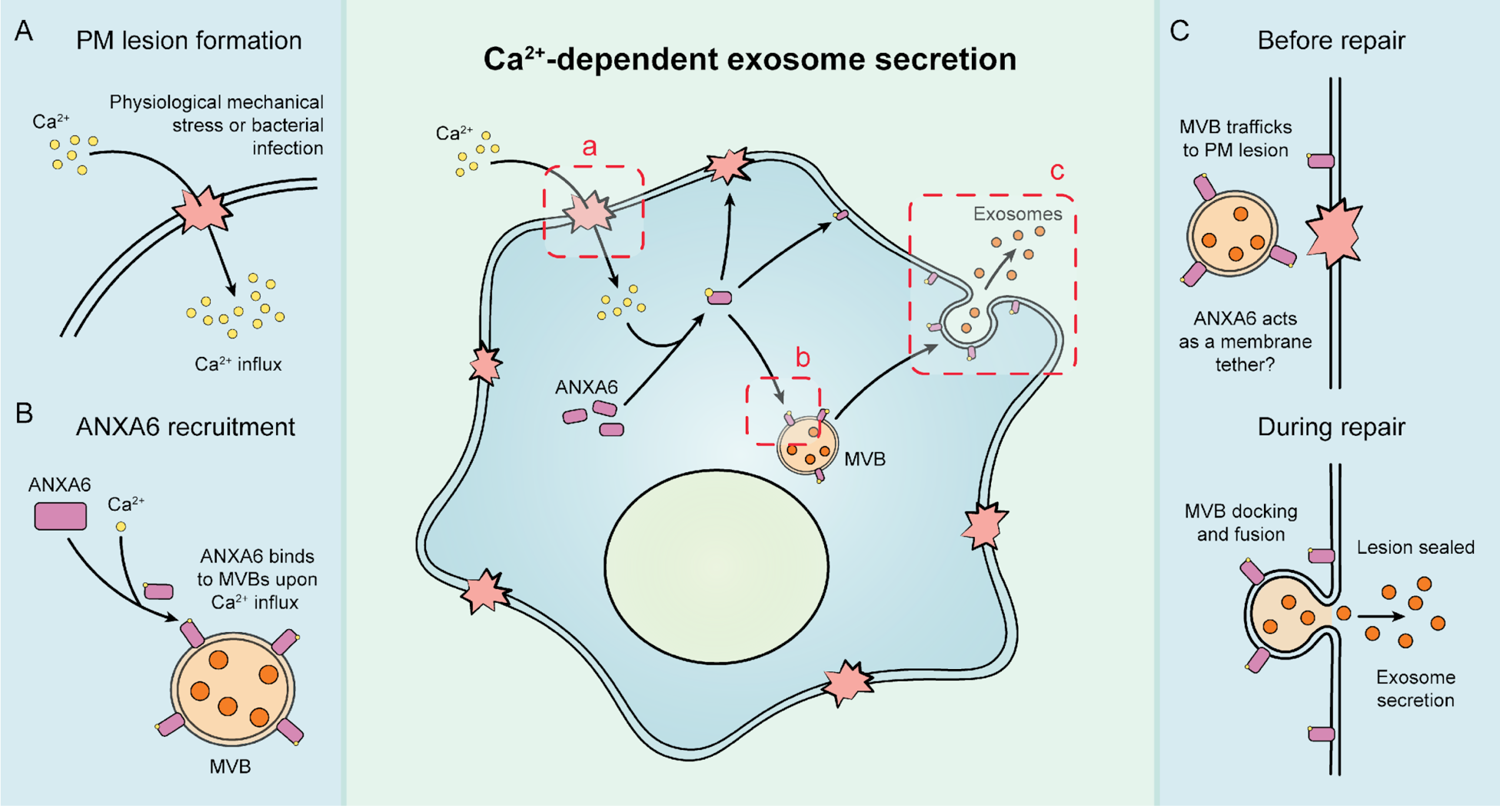
Schematic depicting the current model of Ca^2+^- and ANXA6-dependent MVB exocytosis. (*A*) Upon physiological mechanical stress or bacterial infection, plasma membrane lesions form. This results in the flow of Ca^2+^ from the extracellular space into the cytoplasm. (*B*) This influx of Ca^2+^ mediates the recruitment of ANXA6 to MVBs, which are then transported on microtubules to a plasma membrane lesion. (*C*) The MVBs then dock at the plasma membrane (with ANXA6 serving as a putative membrane tether) and undergo fusion, resulting in plasma membrane repair and exosome secretion.

### Which membrane compartments serve in plasma membrane repair?

We provide evidence that the egress of CD63-positive exosomes from HCT116 cells is coupled to the Ca^2+^-dependent repair of plasma membrane lesions. Microinjection studies in echinoderm eggs were the first to demonstrate that metazoan cells are able to repair wounds to the plasma membrane, and that this repair process is dependent upon the presence of extracellular Ca^2+^ (34–36). Since then, a variety of mechanisms by which metazoan cells are able to repair their plasma membrane (e.g., membrane shedding, organelle exocytosis, clot formation/wound constriction, and membrane internalization) have been proposed (17, 25, 37, 38).

Jaiswal et al. (2002) used total internal reflection fluorescence (TIRF) microscopy to identify the primary membrane compartment responsible for Ca^2+^-dependent exocytosis in non-secretory cells. In their studies, addition of a Ca^2+^ ionophore elicited the fusion of plasma membrane-proximal lysosomes with little or no evidence of involvement of early endosomes, late endosomes/MVBs, or vesicles derived from either the endoplasmic reticulum or Golgi apparatus. Our data that CD63-positive compartments fuse at the plasma membrane upon Ca^2+^ influx is consistent with their findings. However, our results also suggest that endosomes/MVBs can undergo Ca^2+^-dependent exocytosis. We conclude this based on the secretion of MVB cargo vesicles (exosomes) upon Ca^2+^ influx elicited by a Ca^2+^ ionophore, a pore-forming toxin, or physiological levels of mechanical stress. Our results also suggest that both free and pre-docked late endosomes/MVBs can participate in plasma membrane repair as Ca^2+^-dependent exosome secretion induced Ca^2+^ ionophore treatment is microtubule-dependent as indicated by sensitivity to nocodazole. The markers used by Jaiswal et al. (2002) to differentiate between late endosomes (Rab7) and lysosomes (CD63) have now been demonstrated to be present on both membrane compartments (39). The combined data suggest that, in addition to lysosomes, late endosomes/MVBs can undergo Ca^2+^-dependent exocytosis.

Our results are consistent with published studies conducted in sea urchin eggs and cytotoxic T lymphocytes (40, 41). Ca^2+^-dependent plasma membrane repair of sea urchin eggs after micropuncture with glass micropipettes has been demonstrated to be both fusion- and microtubule-dependent as suggested by sensitivity to either botulinum toxins or antibodies that block kinesin-mediated microtubule transport (40, 42–44). Endosomes pre-loaded with Alexa-488-conjugated transferrin traffic towards the plasma membrane upon treatment of cytotoxic T lymphocytes with perforin (41). This result is consistent with our finding that MVBs undergo exocytosis in cells treated with SLO.

Given our findings, we propose that both MVBs and lysosomes undergo Ca^2+^-dependent exocytosis during plasma membrane repair, resulting in the concurrent secretion of exosomes (Fig. 7).

### Role of ANXA6 in plasma membrane repair and exosome secretion

We find that ANXA6 is required for Ca^2+^-dependent fusion of MVBs with the plasma membrane (Fig. 4). Annexin proteins are known to be required for plasma membrane repair (45–50). Depletion of ANXA6 compromises plasma membrane repair, and the N-terminal domain of ANXA6 is insufficient for membrane repair (51, 52). In addition to lysosome exocytosis, plasma membrane repair is accompanied by shedding of damaged membrane at the cell surface (17). Annexins have been invoked in the capping and shedding of plasma membrane lesions (33). Our results extend these conclusions to the fusion of MVBs at the cell surface.

Our data favors a model where ANXA6 is directly involved in tethering an MVB to the plasma membrane during exocytosis. ANXA6 is recruited to CD63-positive compartments in a Ca^2+^-dependent manner (Fig. 3). Additionally, CD63-positive membrane compartments stall near the plasma membrane in Ca^2+^-stimulated, ANXA6-depleted cells (Fig. 4*D*). This conclusion is strengthened by our observation that an anti-ANXA6 antibody blocks exosome secretion in permeabilized cells. Unlike other members of the annexin family, ANXA6 contains two distinct annexin domains, possibly one for each membrane partner of a fusion pair. The N-terminal annexin domain is enriched on the plasma membrane whereas the C-terminal annexin domain is associated with intracellular vesicles (Fig. 6*C*). ANXA6 has also been demonstrated to tether liposomes *in vitro* in the presence of Ca^2+^ (53). Less direct roles in docking/fusion, such as SNARE interactions or actin remodeling as seen for other annexins, remain possible (54, 55).

Although we focused on the role of ANXA6 in Ca^2+^-dependent MVB exocytosis, ANXA2 and CPNE3 were both recruited to MVBs and lysosomes in the presence of calcium, and knockdown of CPNE3 inhibited MVB exocytosis (Fig. 3 and 4*D*). Thus, we hypothesize that these proteins could also be involved in the fusion of MVBs and lysosomes with the plasma membrane during repair.

### EVs in biological fluids

Although our focus has been on exosome secretion, others have reported plasma membrane-derived vesicle secretion stimulated by plasma membrane disruptions (37). Such shedded vesicles may correspond to the low buoyant density vesicles marked by ANXA2 and FLOT2 which we find in the EV fraction produced by ionomycin treatment (Fig. 2*D*).

We demonstrate that SLO-treated or mechanically stressed CD63-Nluc cells secrete exosomes in response to Ca^2+^-dependent membrane repair (Fig. 2*B* and *C*). Such treatments may mimic conditions of physiologic stress *in vivo* and lead to wound-induced EV secretion. This may contribute to the diversity of EVs present in biological fluids, including in samples used for liquid biopsy. Plasma membrane repair is common, especially in certain tissues. In rats, 6.5% of cells in the vascular endothelium of the aorta undergo plasma membrane repair at a given time (56); 20% of muscle cells repair their membrane after muscle contractions (57). Motile cancer cells also have high rates of plasma membrane repair (58). We postulate that exosome secretion during plasma membrane repair may allow for simple detection of tissue-specific damage during routine clinical analysis of blood or urine samples.

The biological functions of EVs released during Ca^2+^-evoked secretion remain unclear. Much attention has focused on a role for EVs in intercellular communication (59). Alternatively, a recent study showed that exosomes act as “decoys” for pore-forming toxins such as α-toxin (60). During staphylococcus infection, cells upregulate exosome production. Instead of binding to the plasma membrane, α-toxin binds to exosomes. Correspondingly, we find that SLO binds exosomes at concentrations that may compete with plasma membrane binding (Fig. S1*A*). Calcium-dependent exosome secretion allows for rapid secretion of high levels of exosomes and may serve as a defense against bacterial toxins that perforate the plasma membrane.

## Materials and Methods

### Key Resources Table

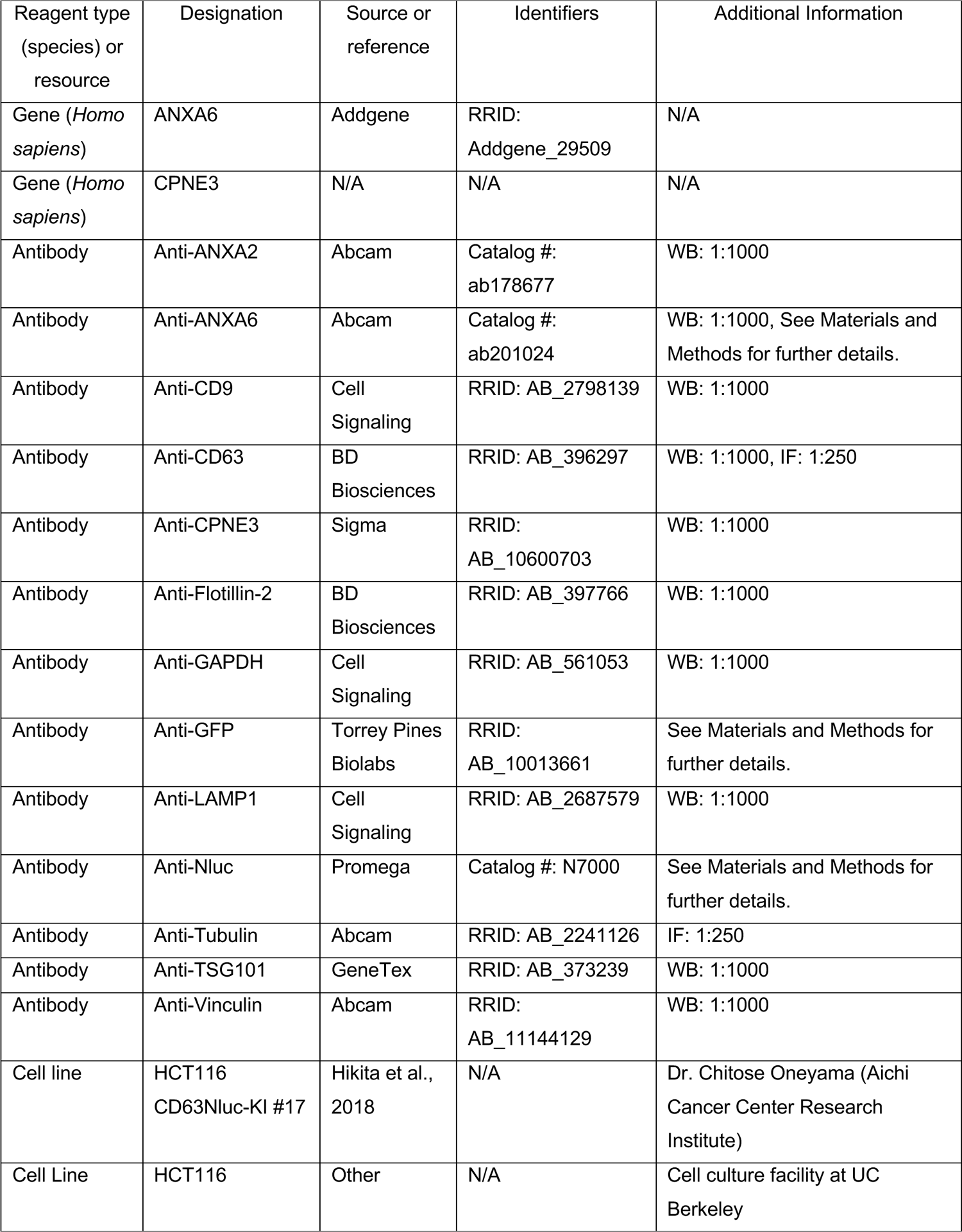
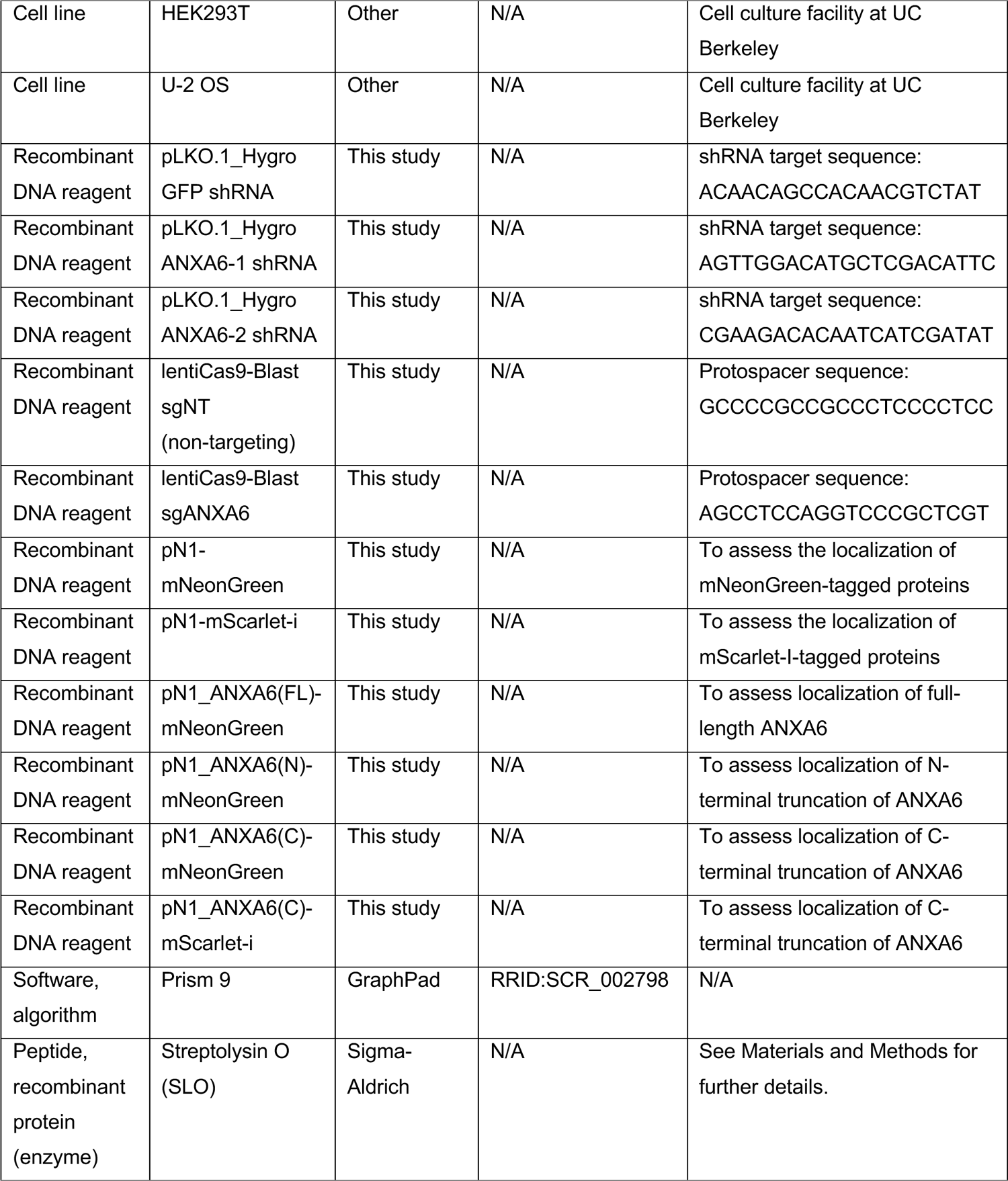

#### Cell lines, media, and general chemicals

HCT116 CD63-Nluc, HEK293T, and U-2 OS cells were cultured at 37°C in 5% CO_2_ and maintained in DMEM supplemented with 10% FBS (Thermo Fisher Scientific, Waltham, MA). Cells were routinely tested (negative) for mycoplasma contamination using the MycoAlert Mycoplasma Detection Kit (Lonza Biosciences). For the experiments detailed in Fig. 2*B* and 2*C*, we cultured HCT116 CD63-Nluc cells in Ca^2+^-free DMEM (Thermo Fisher Scientific). Ionomycin was purchased from Cayman Chemicals. Unless otherwise noted, all other chemicals were purchased from Sigma-Aldrich (St. Louis, MO).

#### Lentivirus production and transduction

HEK293T cells at 40% confluence within a 6-well plate were transfected with 165 ng of pMD2.G, 1.35 µg of psPAX2, and 1.5 µg of a pLKO.1-Hygro or lentiCas9-Blast plasmid using the TransIT-LT1 Transfection Reagent (Mirus Bio) as per the manufacturer’s protocol. At 48 h post-transfection, 1 ml of fresh DMEM supplemented with 10% FBS was added to each well. The lentivirus-containing medium was harvested 72 h post-transfection by filtration through a 0.45 μm polyethersulfone (PES) filter (VWR Sciences). The filtered lentivirus was aliquoted, snap-frozen in liquid nitrogen, and stored at −80°C. For lentiviral transductions, we infected HCT116 CD63-Nluc cells with filtered lentivirus in the presence of 8 μg/ml polybrene for 24 h and the medium was replaced. HCT116 CD63-Nluc cells were selected using 200 μg/ml hygromycin B or 4 μg/ml blasticidin S for 8 d and 6 d, respectively.

#### Immunoblotting

Cells were lysed in phosphate-buffered saline (PBS) containing 1% TX-100 and a protease inhibitor cocktail (1mM 4-aminobenzamidine dihydrochloride, 1 µg/ml antipain dihydrochloride, 1 µg/ml aprotinin, 1 µg/ml leupeptin, 1 µg/ml chymostatin, 1 mM phenylmethylsulfonyl fluoride, 50 µM N-tosyl-L-phenylalanine chloromethyl ketone and 1µg/ml pepstatin) and incubated on ice for 15 min. The whole cell lysate was centrifuged at 15,000xg for 10 min at 4°C and the post-nuclear supernatant was diluted with 6X Laemmli buffer (without DTT) to a 1X final concentration. Samples were heated at 95°C for 5 min and proteins resolved on 4-20% acrylamide Tris-glycine gradient gels (Life Technologies). Proteins were then transferred to polyvinylidene difluoride membranes (EMD Millipore, Darmstadt, Germany), blocked with 5% dry milk in TBS, washed 3X with TBS-T and incubated overnight with primary antibodies in 5% bovine serum albumin in TBS-T. The membranes were then washed again 3X with TBS-T, incubated for 1 h at room temperature with 1:10,000 dilutions of anti-rabbit or anti-mouse secondary antibodies (GE Healthcare Life Sciences, Pittsburgh, PA), washed 3X with TBS-T, washed once with TBS and then detected with ECL-2 or PicoPLUS reagents (Thermo Fisher Scientific) for proteins from cell lysates or EV isolations, respectively.

#### Immunofluorescence and live-cell imaging

For immunofluorescence, we grew cells on coverslips which were then washed once with PBS, fixed in 4% EM-grade paraformaldehyde (Electron Microscopy Science Hatfield, PA) for 15 min at room temperature, washed 3X with PBS and permeabilized/blocked in blocking buffer (2% BSA and 0.02% saponin in PBS) for 30 min at room temperature. Cells were then incubated with a 1:250 dilution of primary antibody and/or phalloidin stain overnight at 4°C, washed 3X with PBS, incubated with a 1:1000 dilution of fluorophore-conjugated secondary antibody for 1 h at room temperature and washed 3X with PBS. Coverslips were then mounted overnight in ProLong-Gold antifade mountant with DAPI (Thermo Fisher Scientific) and sealed with clear nail polish before imaging. Images were acquired on an Echo Revolve Microscope using the 60X Apo Oil Phase, NA 1.42 objective.

For live-cell imaging, we plated cells on 35mm glass-bottom imaging dishes (MatTek) for 16 h followed by transfection with the intended plasmids using Lipofectamine 2000 as per the manufacturer’s protocol. The medium containing the transfection mixture was removed ∼5 h post-transfection and replaced with DMEM supplemented with 10% FBS. At 16-18 h post-transfection, the medium was replaced with Fluorobrite DMEM (Gibco) supplemented with 1X GlutaMAX (Gibco) prior to imaging. Images were acquired using an LSM900 confocal microscope system (ZEISS) within a temperature-controlled environment chamber (37°C and 5% CO_2_) using Airyscan 2 mode and a 63X Plan-Apochromat, NA 1.40 objective. The concentrations utilized for each plasmid were as follows: pN1-ANXA6(FL)-mNeonGreen (200 ng/ml), pN1-ANXA6(N)-mNeonGreen (100 ng/ml), pN1-ANXA6(C)-mNeonGreen (100 ng/ml), and pN1-ANXA6(C)-mScarlet (100 ng/ml).

#### EV isolation and fractionation by iodixanol buoyant density gradient equilibration

Fresh aliquots of 5%, 7.5% 10%, 12.5%, 15%, 17.5% 20%, 22.5%, and 25% (v/v) iodixanol solutions were prepared by mixing appropriate volumes of Solution B (0.25 M sucrose, 2 mM MgCl_2_, 1 mM EDTA, 20 mM Tricine-NaOH, pH 7.8) and Solution D (41.7 mM sucrose, 2 mM MgCl_2_, 1 mM EDTA, 20 mM Tricine-NaOH, pH 7.8, 50% (w/v) iodixanol). Iodixanol gradients were prepared by sequential 500 µl overlays of each iodixanol solution in a 5 ml SW55 tube, starting with the 25% iodixanol solution and finishing with the 5% iodixanol solution. After the addition of each iodixanol solution, the SW55 tube was flash frozen in liquid nitrogen. Complete iodixanol gradients were stored at −20°C and thawed at room temperature for 45 min prior to use.

For the iodixanol gradients detailed in Fig. 2*D*, conditioned medium (240 ml) was harvested from vehicle- or ionomycin-treated HCT116 CD63-Nluc cells. All subsequent manipulations were completed at 4°C. Cells and large debris were removed by low-speed sedimentation at 1,000xg for 15 min in a Sorvall R6+ centrifuge (Thermo Fisher Scientific) followed by medium-speed sedimentation at 10,000xg for 15 min using a fixed angle FIBERlite F14-6 x 500 y rotor (Thermo Fisher Scientific). The supernatant fraction was then centrifuged at 32,000 RPM for 1.25 h in a SW32 rotor. The high-speed pellet fractions were resuspended in PBS, pooled, loaded at the top of a prepared iodixanol gradient, and centrifuged in a SW55 rotor at 36,500 RPM for 16 h with minimum acceleration and no brake. Fractions (200 µl) were collected from top to bottom. An aliquot of each fraction was saved for luminescence analysis and the rest was diluted in 6X Laemmli buffer (without DTT) for immunoblot analysis. Density measurements were taken using a refractometer.

For the iodixanol gradients detailed in Fig. 2*E*, we harvested conditioned medium from vehicle- or ionomycin-treated HCT116 CD63-Nluc cells grown in a 12-well plate. All subsequent manipulations were completed at 4°C. Cells and large debris were removed by low-speed sedimentation at 1,000xg for 15 min followed by medium-speed sedimentation at 10,000xg for 15 min in an Eppendorf 5430 R centrifuge (Eppendorf, Hamburg, Germany). Aliquots (200 µl) of conditioned medium from the supernatant of the medium-speed centrifugation were loaded at the top of a prepared iodixanol gradient and centrifuged in a SW55 rotor at 36,500 RPM for 16 h with minimum acceleration and no brake. Fractions (200 µl) were collected from top to bottom and analyzed for luminescence. Density measurements were taken using a refractometer.

#### Immunoisolation of MVBs and Ca^2+^-dependent binding proteins

HCT116 CD63-Nluc cells were grown to ∼90% confluence in 7 x 150 mm dishes. Cells were scraped into 5 ml of cold PBS per plate and centrifuged at 200xg for 5 min at 4°C. The cold PBS was aspirated, and the cell pellet was resuspended in 2 volumes of cold lysis buffer (136 mM KCl, 10 mM KH_2_PO_4_ pH 7.4, 1 mM DTT, protease inhibitor cocktail [see Immunoblotting section] and 6% Optiprep (w/v)). The cell slurry was passed 14 times through a 25-gauge syringe in a cold room, and the post-nuclear supernatant was prepared by centrifugation of lysed cells at 1000xg for 10 min at 4°C. The post-nuclear supernatant was diluted 1:2 with lysis buffer.

Beads from 300 µl of magnetic Protein G Dynabeads (Thermo Fisher Scientific) slurry were sedimented with a magnetic tube rack and resuspended in lysis buffer. The bead slurry was split evenly into 3 tubes and then re-centrifuged. The diluted post-nuclear supernatant was divided between the 3 tubes. Tube #1 also received 1 mM CaCl_2_ (Beads only control), tube #2 also received 1 mM CaCl_2_ and 5 µg anti-Nluc antibody (Ca^2+^-treated), and tube #3 also received 1 mM EGTA and 5 µg anti-Nluc antibody (EGTA treated). The reaction was incubated for 15 min at room temperature. Beads were washed 3 times with lysis buffer. The beads only control and Ca^2+^-treated samples both had 2 mM CaCl_2_ added to the wash buffer. The EGTA-treated sample was washed with lysis buffer containing 2 mM EGTA. The beads only control and Ca^2+^-treated samples were eluted with 50 µl lysis buffer containing 2 mM EGTA and the EGTA-treated sample was eluted with lysis buffer containing 2 mM CaCl_2_. All three samples were eluted again with 50 µl lysis buffer containing 0.2% TX-100.

#### CD63-Nluc exosome secretion assay

HCT116 CD63-Nluc cells were grown to ∼80% confluence in 24-well plates. All subsequent manipulations were performed at 4°C. Conditioned medium (200 µl) was taken from the appropriate wells, added to a microcentrifuge tube and centrifuged at 1,000xg for 15 min in an Eppendorf 5430 R centrifuge (Eppendorf, Hamburg, Germany) to remove intact cells. Supernatant fractions (150 µl) from the low-speed sedimentation were moved to a new microcentrifuge tube and centrifuged at 10,000xg for 15 min to remove cellular debris. Supernatant fractions (50 µl) from this medium-speed centrifugation were then utilized to measure CD63-Nluc exosome luminescence. During these centrifugation steps, the cells were placed on ice, washed once with cold PBS and lysed in 200 µl of PBS containing 1% TX-100 and protease inhibitor cocktail.

To measure CD63-Nluc exosome secretion, we prepared a master mix containing the membrane-permeable Nluc substrate and a membrane-impermeable Nluc inhibitor using a 1:1000 dilution of Extracellular NanoLuc Inhibitor and a 1:333 dilution of NanoBRET^TM^ Nano-Glo Substrate into PBS (Promega, Madison, WI). Aliquots of the Nluc substrate/inhibitor master mix (100 µl) were added to 50 µl of the supernatant fraction from the medium-speed centrifugation, vortexed briefly and luminescence was measured using a Promega GlowMax 20/20 Luminometer (Promega, Madison, WI). An aliquot (1.5 µl) of 10% TX-100 was then added to each reaction tube for a final concentration of 0.1% TX-100 and the sample was briefly vortexed before luminescence was measured again. For the intracellular normalization measurement, the luminescence of 50 µl of cell lysate was measured using the Nano-Glo Luciferase Assay kit (Promega, Madison, WI) as per the manufacturer’s protocol. The exosome production index (EPI) for each sample is calculated as follows: EPI = [(Medium) – (Medium + 0.1% TX-100)] / Cell Lysate

#### Mechanical stress experiments

Two 15 cm plates of CD63-Nluc cells were harvested with 10 ml of Accutase and diluted with 40 ml of Ca^2+^-free DMEM. A cell slurry (16 ml) was added to three tubes and centrifuged at 300xg for 5 min at room temperature. Cells were gently resuspended in either 0.5 ml Ca^2+^-free DMEM or Ca^2+^-free DMEM + 2 mM Ca^2+^ (final). Cells were pumped through a 30-gauge needle at a flow rate of 3.5 µl/sec (τ = 4Qη/πR^3^ = ∼89 dyn/cm^2^, where Q = 0.0035 cm^3^/s, η = 0.01 dyn*s/cm^2^, 30 G average internal radius = 7.94×10^-3^ cm) or twice the flow rate of 7 µl/sec (τ = 4Qη/πR^3^ = 178 dyn/cm^2^, where Q = 0.0035 cm^3^/s, η = 0.01 dyn*s/cm^2^, 30 G average internal radius = 7.94×10^-3^ cm) using a Harvard Apparatus syringe pump (Catalog No. 98-4730). Cells were incubated for 5 min at 37°C before being placed back on ice. The cell suspension was centrifuged at 300xg for 5 min and the supernatant fraction was then filtered through a 0.45 µm PES filter. Exosome secretion was measured as described in paragraph 2 of “CD63-Nluc exosome secretion assay,” in this assay without a cell lysate measurement.

#### Isolation of cytosol from cultured human cells

HCT116 WT cells were grown to ∼90% confluence in 20 x 150 mm dishes. All subsequent manipulations were performed at 4°C. Each 150 mm dish was washed once with 10 ml of cold PBS and then harvested by scraping into 5 ml of cold PBS. The 5 ml cell suspension was then used to harvest cells from 4 additional 150 mm dishes and this process was repeated until all the cells were harvested. The cells were then collected by centrifugation at 200xg for 5 min. The supernatant fraction was discarded, and the cell pellet was resuspended in 3 ml of cold hypotonic lysis buffer (20 mM HEPES, pH 7.4, 10 mM KCl, 1 mM EGTA, 1 mM DTT, and protease inhibitor cocktail) and placed on ice. After 15 min, the cell suspension was transferred to a pre-chilled 7 ml Dounce homogenizer, and the cells were mechanically lysed by ∼80 strokes with a tight-fitting Dounce pestle. The lysed cells were centrifuged at 1,000xg for 15 min to sediment intact cells and nuclei. The post-nuclear supernatant was then centrifuged at 32,500 RPM (∼128,000xg) for 30 min in an Optima XE-90 ultracentrifuge (Beckman Coulter). The supernatant (cytosol fraction) was collected conservatively without disturbing the pellet and then concentrated using a 4 ml Amicon®-3k concentrator. The final concentration of the cytosol fraction was ∼40 mg/ml. The cytosol was distributed in aliquots, snap-frozen in liquid nitrogen, and stored at –80°C until use. The rat liver cytosol was prepared as described in (61).

#### Cell-free reconstitution of CD63-Nluc exosome secretion

SLO was pre-activated in PBS containing 10 mM DTT at 37°C for 2 h, distributed in aliquots into low-retention microcentrifuge tubes, snap-frozen in liquid nitrogen and stored at –80°C until use. The protein concentration of each SLO batch was determined by a Bradford assay.

HCT116 CD63-Nluc cells were grown to ∼80% confluence in 24-well Poly-D-Lysine (PDL) coated plates. The PDL plate was placed on ice and the cells were washed once with PBS containing 1 mM EGTA. The PBS wash was aspirated, replaced with 200 µl of cold transport buffer (20 mM HEPES, pH 7.4, 250 mM D-sorbitol, 120 mM KCl, 10 mM NaCl, 2 mM MgCl_2_, 1.2 mM KH_2_PO_4_, 1 mM EGTA, and protease inhibitor cocktail) supplemented with 0.6 µg/ml of pre-activated SLO and incubated at 4°C for 15 min. Unbound SLO was aspirated, and the cells were washed once with cold transport buffer. The cells were then permeabilized by the addition of pre-warmed transport buffer containing 2 mM DTT followed by a 10 min incubation at 37°C. The permeabilized CD63-Nluc cells were then washed at 4°C in transport buffer, high-salt transport buffer (containing 1 M KOAc) and finally transport buffer (10 min each wash) to deplete cytosol.

Complete cell-free exosome secretion assays (200 µl) consisted of permeabilized CD63-Nluc cells, cytosol (4 mg/ml final concentration), 20 µl 10X ATP^r^ (10 mM ATP, 400 mM creatine phosphate, 2 mg/ml creatine phosphokinase, 20 mM HEPES, pH 7.2, 250 mM D-sorbitol, 150 mM KOAc, 5 mM MgOAc), 3 µl of 10 mM GTP, 4 µl of 100 mM CaCl_2_ (2 mM final concentration) and cold transport buffer. The concentration of blocking IgG antibodies utilized in Fig. 5*E* was 25 µg/ml. The assembled reaction mixes were added to the permeabilized CD63-Nluc cells for 5 min on ice prior to placing the entire 24-well PDL coated plate in a 30°C water bath for 2 min to stimulate exosome secretion. The cells were then placed back on ice, and 100 µl of each reaction supernatant was loaded into a 0.4 µm AcroPrep filter plate (Pall Corporation) and centrifuged at 1,500xg for 1 min in an Eppendorf 5810 R centrifuge (Eppendorf, Hamburg, Germany) to collect CD63-Nluc exosomes. During this centrifugation step, 100 µl of cold transport buffer containing 2% TX-100 and protease inhibitor cocktail was added to each well of the 24-well PDL coated plate to lyse the cells and bring the volume up to 200 µl. Filtrate aliquots (50 µl) and 50 µl of the cell lysate were used to measure exosome secretion and normalize to the number of cells per reaction, respectively, using the cell-based CD63-Nluc exosome secretion assay protocol detailed above.

For the cell-free exosome secretion assay detailed in Fig. 5*D*, the nucleotide-depleted rat liver cytosol was generated by using a HiTrap Sephadex G-25 Desalting Column (Cytiva Life Sciences) as per the manufacturer’s protocol.

## Additional Information

### Funding

**Funder** Howard Hughes Medical Institute, Sergey Brin Family, Foundation

**Grant Reference Number** Investigator, Scientific Director, ASAP

**Author** Randy Schekman, Randy Schekman

The funders had no role in study design, data collection and interpretation, or the decision to submit the work for publication.

### Author Contributions

J.K.W., J.M.N., and R.S. designed research; J.K.W. and J.M.N. performed research; J.K.W., J.M.N., and R.S. analyzed data; and J.K.W., J.M.N., and R.S. wrote the paper.

## Acknowledgements

We dedicate this work to Bob Lesch, our lab manager for the past several decades who was tragically taken from us by an accident this past year. We thank Criss Hartzell for reading and providing helpful comments on this manuscript. We also thank Isabelle Lehman for experimental assistance. Additionally, we would also like to thank the staff at the UC Berkeley shared facilities, the Cell Culture Facility (Alison Killilea), the Vincent J Coates Proteomics Facility (Lori Kohlstaedt) and the DNA Sequencing Facility. J.M.N is supported by a National Science Foundation Graduate Research Fellowship. R.S. is an Investigator of the Howard Hughes Medical Institute, a Senior Fellow of the UC Berkeley Miller Institute of Science, and Scientific Director of Aligning Science Across Parkinson’s Disease (ASAP).

---

**Fig. S1.**
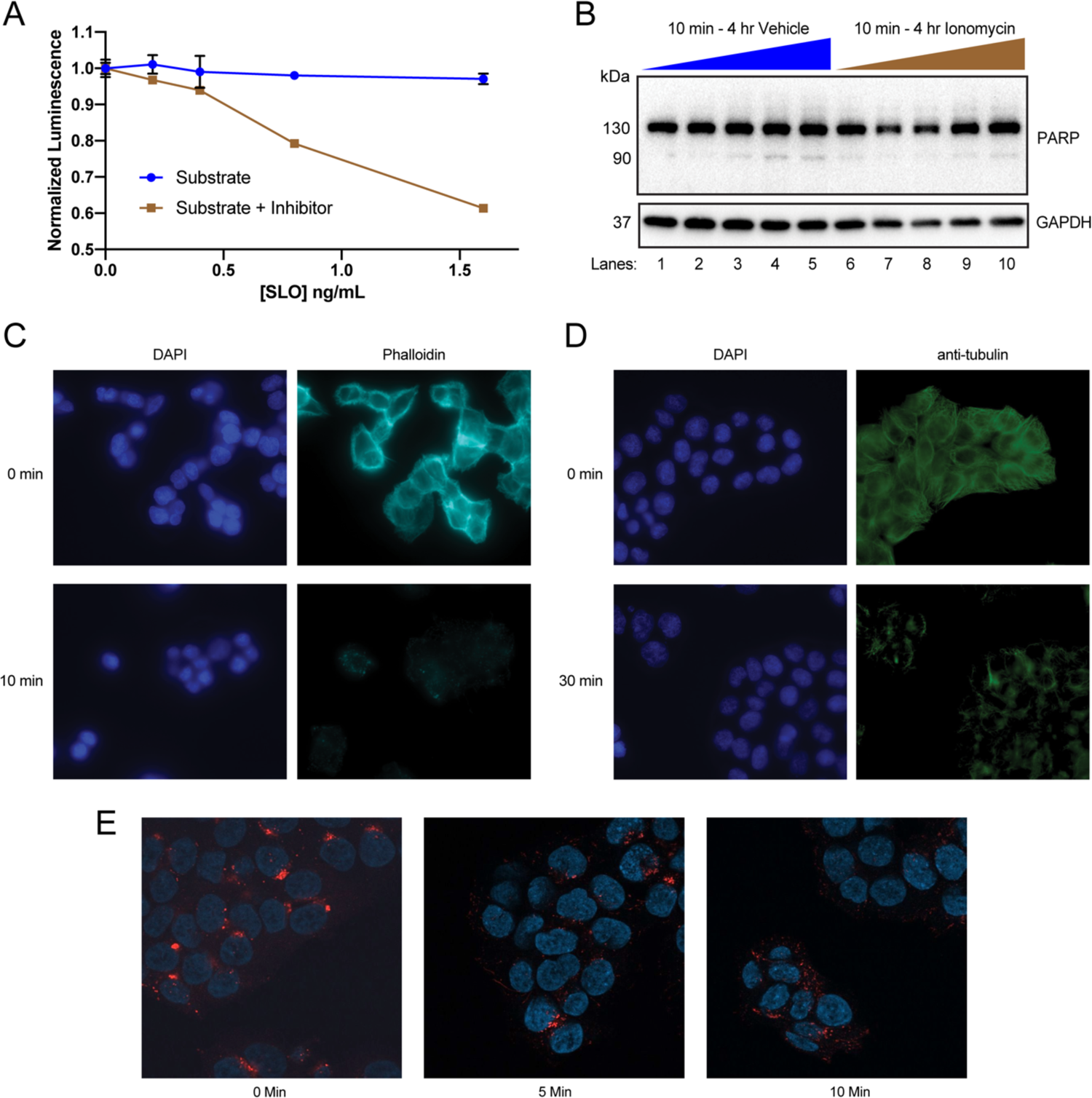
(*A*) Normalized luminescence derived from the 10k x g conditioned medium supernatant from CD63-Nluc cells with or without the addition of the membrane-impermeable Nluc inhibitor, as a function of increasing SLO concentrations, is shown. (*B*) PARP and GAPDH immunoblots from HCT116 CD63-Nluc cells during a 4 h ionomycin (5 µM) time course. (*C*) Phalloidin staining after treatment with 1 µM latrunculin A or (*D*) tubulin immunofluorescence after treatment with 10 µM nocodazole. Data plotted represent the means from 2 independent experiments and error bars represent standard deviations. (*E*) LAMP1 (Red) immunofluorescence over 10 min of 5 µM ionomycin is shown. DAPI staining is shown in blue.

**Fig. S2.**
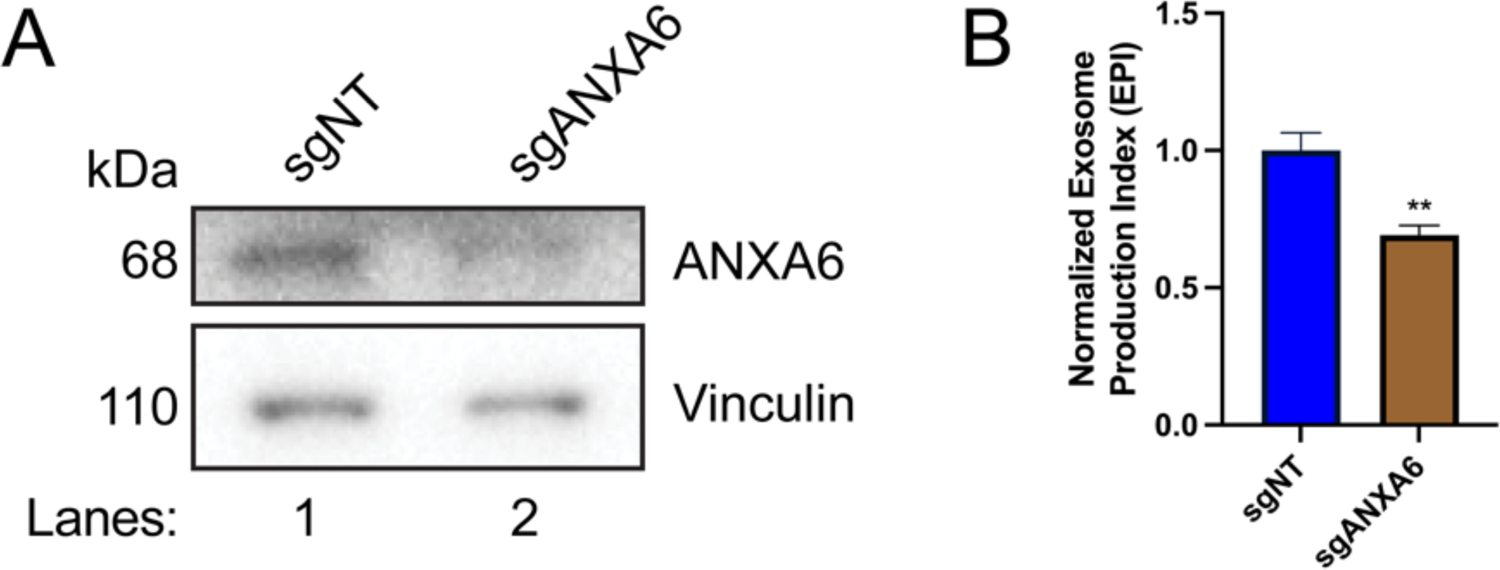
Exosome secretion from polyclonal ANXA6 KO cells. (*A*) Immunoblot analysis of ANXA6 and vinculin expression from sgNT and sgANXA6 CD63-Nluc cells are shown. (*B*) Normalized exosome production derived from sgNT and sgANXA6 CD63-Nluc cells treated with 5 µM ionomycin for 30 min are shown. Data plotted represent the means from 3 independent experiments and error bars represent standard deviations. Statistical significance was performed using a Student’s T-test (**p<0.01).

